# Memory-safe high-performance sequence mapping with rammap

**DOI:** 10.64898/2026.05.26.726289

**Authors:** Jeremy R. Wang, Heng Li

**Affiliations:** Department of Pathology and Laboratory, Medicine Department of Genetics, University of North Carolina at Chapel Hill; Department of Data Science, Dana-Farber Cancer Institute, Department of Biomedical Informatics, Harvard Medical School

## Abstract

We introduce a reimplementation of the widely used mapping tool minimap2 in Rust called `rammap`. We demonstrate perfect concordance with minimap2, enabling its backwards compatibility as a drop-in replacement for minimap2-based workflows. Additionally, rammap implements performance optimizations for modern architectures and applications, including AVX512 and WASM v128 SIMD support for dynamic programming alignment and SIMD-accelerated chaining. These achieve comparable or better performance than minimap2 across diverse mapping workloads while maintaining Rust’s stronger memory safety constraints. The rammap API exposes both SIMD-accelerated sequence-sequence alignment modules and full mapping pipelines for use as an integrated library. Lastly, we describe the modular architecture and provide examples illustrating the extensibility of major mapping components, including seeding, chaining, and gap-filling/extension to support development of improved or domain-specific mapping components within a high-performance framework.

## Introduction

Read mapping is the cornerstone of modern high-throughput sequencing data analysis. The seed-and-extend paradigm has long underpinned efficient read mapping in widely used tools such as BLAST [1], bwa [2], and bowtie2 [3]; its seed-chain-extend extension, in which colinear chain assembly across multiple anchors is performed as an explicit intermediate stage, is exemplified by BLAT [4], BWA-MEM [5] and its SIMD(Single Instruction, Multiple Data)-accelerated successor BWA-MEM2 [6], and minimap2 [7]. Since its introduction in 2018, minimap2 has been the *de facto* standard for mapping of long read single-molecule (“third-generation”) sequencing data due to its efficient implementation and minimizer-based seeding scheme that takes advantage of anchor arrangement in long error-prone sequences. Minimap2 alignments serve as the requisite input to many widely used bioinformatics tools and workflows, which are often sensitive to the particular algorithmic choices, sensitivities, and parameter settings thereof. It is a critical part of the growing long-read bioinformatic infrastructure. However, minimap2′s low-level C implementation, while performant, requires careful manual memory management and is challenging to both extend and integrate into modern bioinformatics tools.

We introduce rammap, representing a complete reimplementation of minimap2′s mapping approach in Rust. We demonstrate complete backwards compatibility and comparable memory and CPU performance across various mapping tasks, making rammap a viable drop-in replacement for minimap2 with improved library API, wider SIMD support, extensible framework, and Rust’s stricter memory safety architecture.

## Background

Modern seed-chain-extend mapping frameworks apply several algorithmic and computational approaches to enable efficient read-to-reference, read-to-read, and reference-to-reference mapping for data generated by extant high-throughput sequencing technologies. These technologies fall into several broad classes that place qualitatively different demands on a mapping tool. Short-read sequencing-by-synthesis platforms (ex. Illumina [8], BGI/MGI, Element) produce 100–300 bp reads at very high per-base accuracy and extreme throughput, prioritizing seed lookup and small-footprint output. Single-molecule long-read platforms—Pacific Biosciences (PacBio) and Oxford Nanopore Technologies (ONT)—produce reads ranging from kilobases to megabases at lower but steadily improving accuracy [9], with PacBio HiFi reads achieving short-read-comparable accuracy at multi-kilobase length [10]; their length and error structure rewards chaining over many anchors per read and benefit disproportionately from chain-stage acceleration. Beyond standard genomic resequencing, several application domains impose additional structural requirements: chromosomal conformation capture (Hi-C [11]) produces paired reads spanning long-range contacts and requires that mate pairs be mapped independently across distant genomic loci; RNA and cDNA sequencing requires splice-aware alignment to bridge introns; and assembly-to-reference and read-overlap mapping place still different load profiles on the chaining and extension stages. A general-purpose mapper will ideally handle each of these regimes efficiently within a single computational framework.

Two prior efforts have sought to extend or repackage minimap2 in directions adjacent to the present work. Mm2-fast [12] is a fork of minimap2 that introduces hardware-specific accelerations for x86 platforms, including AVX512 support for vectorized chaining, vectorized base-level dynamic programming (DP), and a learned-index seed lookup, achieving a reported 1.8-fold end-to-end speedup over upstream minimap2 while preserving identical output. However, it remains an x86 hardware-specific C implementation and algorithmic changes in the seed lookup step mean that to achieve the full speedup, the output explicitly differs from minimap2. The specific optimization for speed also results in a significant increase in memory usage (∼2x for the human genome). Performance gains— including the reported 1.8x speedup—are focused on long reads with high error rates and, as such, are less significant as modern long read error rates have gone down. Minimap2-rs [13] is a thin Rust wrapper over minimap2, making minimap2 callable from Rust by binding to the existing C library rather than reimplementing the alignment kernels themselves. While this approach explicitly makes minimap2 accessible to Rust applications, it does not extend Rust’s safety guarantees into the underlying alignment kernels nor support native Rust integration and extension of the independent mapping stages.

In this work, we provide a fully Rust-native implementation of the mapping strategy underlying minimap2, with complete backwards compatibility. This approach permits both novel implementation of higher bandwidth (AVX512) SIMD-accelerated chaining and alignment similar to mm2-fast, and an accessible native Rust API to support direct integration into modern bioinformatics tooling.

## Methods

By design, the mapping approach, program architecture, and specific algorithmic decisions largely match minimap2 [7]. However, we implement several notable features to improve performance across newer CPU architectures and to enable broader applications and extensibility.

### Minimizer sketching and indexing

Rammap sketches the reference using the (*k, w*)-minimizer scheme of Roberts et al. [14], in which each window of *w* consecutive *k*-mers contributes its hash-minimized *k*-mer as an indexed anchor. The minimap2 binary index format is supported as both an input and an output, allowing previously built minimap2 indices to be reused without reconstruction. Index construction is structured to bound peak memory rather than to maximize build-time concurrency: reference contigs are sketched sequentially, with the underlying sequence for each freed before the next is read, so the resident set during indexing reflects only the growing minimizer buckets and the 4-bit-packed reference rather than all input sequences held simultaneously. Sketched minimizers are distributed across a fixed set of buckets that are then sorted in parallel and consolidated into per-bucket open-addressing hash tables, with per-bucket consolidation performed sequentially to limit peak memory allocation. The bucketed layout also permits lock-free lookup against a shared flat positions array with cache-friendly bucket-local probing.

### Chaining

Colinear seed chains are constructed via a DP formulation that scores predecessors by anchor distance, allowable gap, and a logarithmic gap-cost term. The standard *O*(*nh*) chaining DP, with bounded predecessor window *h*, is used for read-mapping presets and is exposed through SIMD-vectorized inner loops that score multiple predecessor anchors in parallel. For long-range chaining used by the assembly-to-reference presets, rammap implements an RMQ-tree–accelerated *O*(*n* log *n*) variant that mirrors minimap2′s implementation.

### Gap filling and extension

Adjacent anchors within a chain are joined by base-level alignment using one of several dynamic programming kernels, including single-affine, dual-affine, splice-aware gap-cost variants, and SIMD-accelerated variants of each. Rammap supports a broad range of SIMD instruction sets across different platform architectures, including SSE2, SSE4.1, AVX2, and AVX512BW (x86_64), NEON (aarch64), and SIMD128 (WebAssembly), with separate kernels for both alignment DP and chaining stages. High-bandwidth AVX512 instructions provide clear speedups, as first demonstrated by mm2-fast [12], especially evident in DP-heavy workloads (ex. map-hifi, map-ont).

### Memory safety

Rust’s compiler enforces memory safety statically through analysis of object lifetimes and borrowing, ruling out, by construction, the classes of memory error—use-after-free, buffer overflow, data race —that have historically plagued C-language tools. The SIMD load and store instructions used by rammap’s vectorized kernels lie outside the scope of the compiler’s static analysis, and rammap’s SIMD kernels therefore necessarily contain explicitly memory-unsafe operations confined to those kernels. For applications in which provable memory safety across the entire pipeline is required, rammap can be configured to disable all SIMD code paths and fall back to a fully scalar implementation; this scalar path is necessarily slower than the SIMD-accelerated path but produces identical output and contains no explicitly memory-unsafe operations. A number of rammap’s Rust dependencies include strictly unsafe code, including for memory allocation, gzip handling, and rayon parallelism - these are very widely used and stable libraries that carry minimal risk in practice.

In the process of developing rammap, we identified and fixed two previously unknown bugs in minimap2 that led to unintentional behavior. One involved accessing memory outside the DP band (but within the allocated heap), and the other modifying an active vector in place. Both classes of unintended memory access behavior are exposed and explicitly handled in rammap, further emphasizing the benefits of Rust’s memory safety architecture in this context. These have been fixed in minimap2 v2.31.

### WebAssembly support

Rammap can be compiled directly to native WebAssembly, using WebAssembly’s 128-bit SIMD extension for the dynamic-programming and chaining kernels. WebAssembly additionally supports multiprocessing across browser threads via shared memory, enabling efficient mapping entirely inside the browser.

### Library and API

Beyond its command-line interface, rammap is structured as a library that exposes both the full mapping pipeline and the underlying base-level alignment routines to other applications. Index construction is supported from a reference FASTA, from a pre-built minimap2 binary index file, or from sequences held in memory; the same preset-derived parameter sets used by the command-line interface are used by the library, ensuring identical mapping behavior. Mapping is invoked per query and the returned result exposes hit-level information—chain anchors, CIGAR operations, mapping coordinates, and auxiliary tag data—as native data structures. Library instances are safe to share across worker threads without external synchronization. The library additionally exposes the lower-level DP routines that perform single-pair sequence alignment, providing a performant interface for callers that need SIMD-accelerated nucleotide-level alignment without a full mapping pipeline. The library is intended to occupy a similar role to minimap2-rs [13], but as a native reimplementation rather than a binding to an external C library, extending Rust’s memory-safety guarantees uniformly across the alignment kernels.

### Extensibility

Each major stage of the rammap pipeline—seed selection, chaining, and base-level extension—is defined behind a stage-level interface, with a default production implementation that may be substituted without modifying the rest of the pipeline. We include example alternative implementations for each major mapping stage, including syncmer-based [15] and randstrobe-based [16] seed selectors; DP-based, RMQ-tree-based, and a simple greedy chainer; and an unoptimized scalar Needleman–Wunsch [17] alignment extension implementation that requires no SIMD support. Each implementation follows the common API contract, making them technically interchangeable. We anticipate this structure facilitating the development of domain-specific seeding, chaining, or alignment strategies—for instance, sketching schemes tuned to high-error long reads, chain-cost models adapted to specific transcriptomic or structural-variant contexts, or alignment kernels exposing other hardware-specific accelerators— within the rammap architecture.

## Results

Rammap and minimap2 (2.30-r1299) performance was compared on a high performance server blade with Intel Xeon Gold 6140 CPU (2.3 Ghz, supporting AVX512), allocated 8 threads and 32GB RAM. Publicly available ONT, PacBio (HiFi and Onso), Illumina (PCR-free and Hi-C), BGISEQ, and Element AVITI DNA sequencing data and ONT RNA-sequencing data (Table 1) were evaluated across various presets and alignment modes. Datasets are drawn from the Genome in a Bottle (GIAB) reference releases [18] and the Human Pangenome Reference Consortium (HPRC) production data releases [19], supplemented by a public direct-RNA reference run for ONT’s RNA004 chemistry. We compare CPU time, wall time, and peak resident memory at eight threads for both tools to account for different realistic SIMD and multithreading efficiency across both tools (Table 2).

**Table 1:**
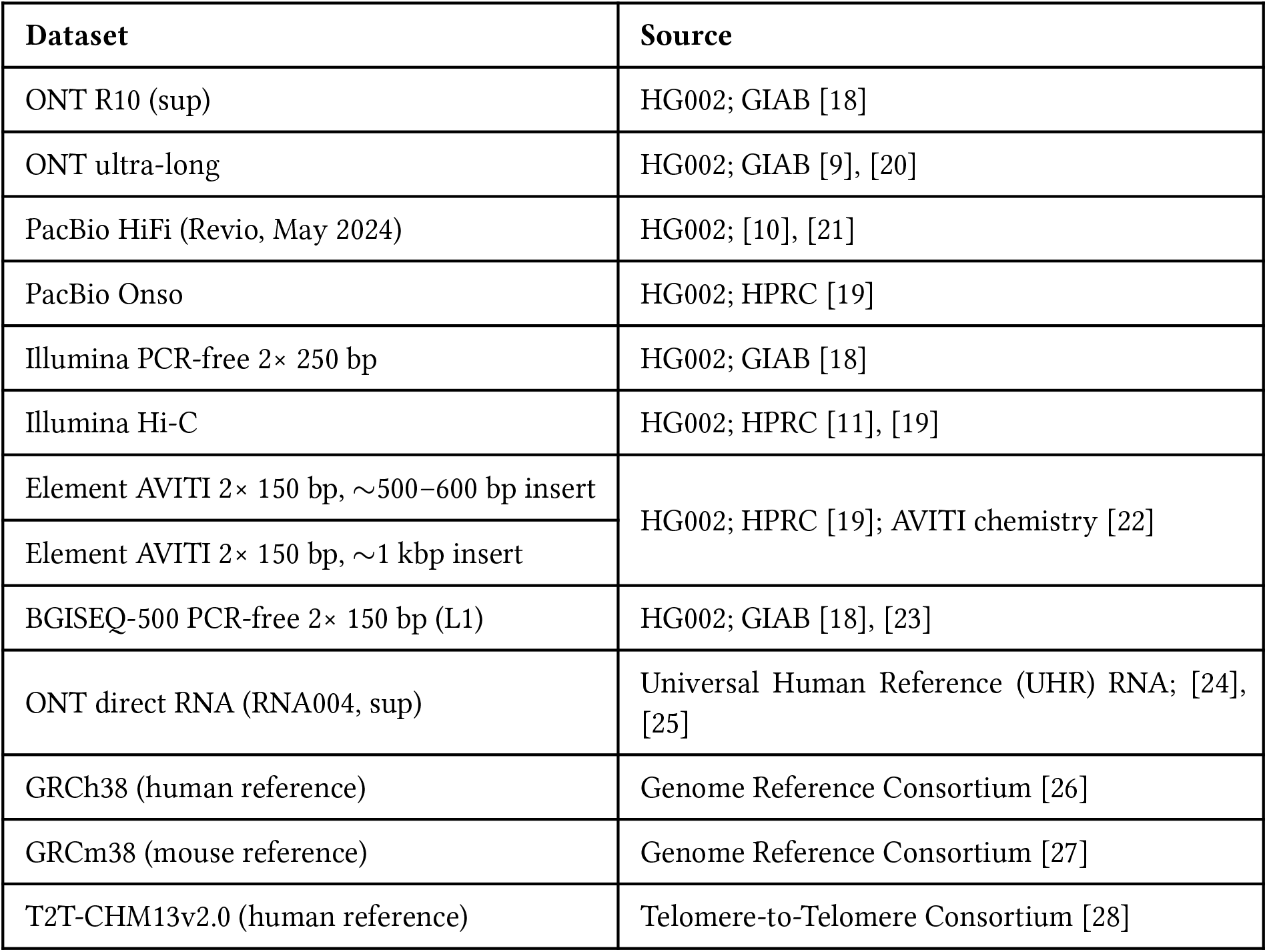
Datasets used for performance benchmarks. Performance characteristics across these datasets are summarized below. In all cases, rammap produces identical alignment output to minimap2 (with the exception of SAM headers including the tool name itself).

**Table 2:**
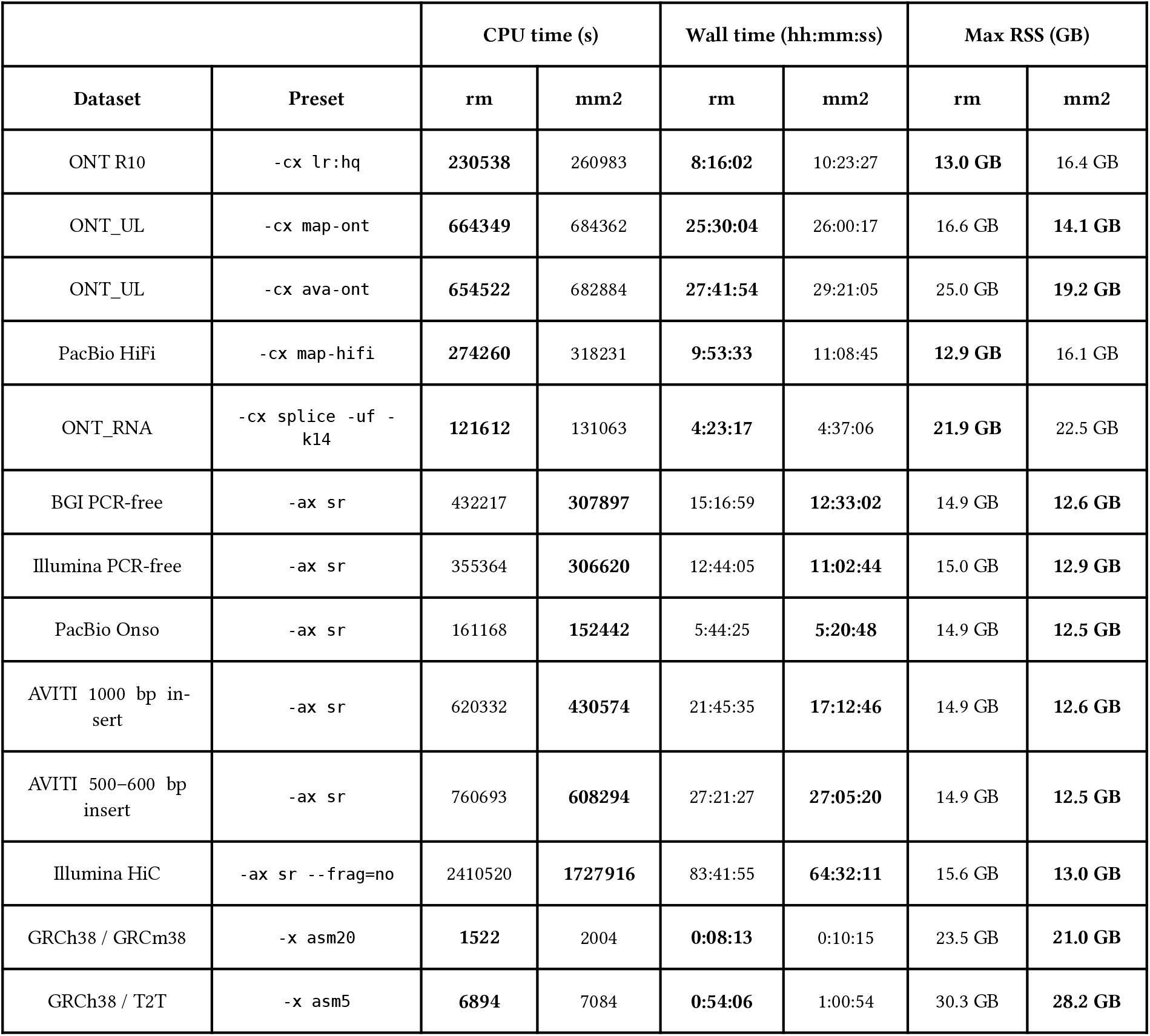
Performance on x86_64 with AVX512 (Intel Xeon Gold 6140); 8 threads, 32 GB RAM. **Bold** marks the better outcome per metric (lower is better). All show identical output between minimap2 and rammap.

|  |  | CPU time (s) |  | Wall time (hh:mm:ss) |  | Max RSS (GB) |  |
| --- | --- | --- | --- | --- | --- | --- | --- |
| Dataset | Preset | rm | mm2 | rm | mm2 | rm | mm2 |
| ONT R10 | -cx lr:hq | **230538** | 260983 | **8:16:02** | 10:23:27 | **13.0 GB** | 16.4 GB |
| ONT_UL | -cx map-ont | **664349** | 684362 | **25:30:04** | 26:00:17 | 16.6 GB | **14.1 GB** |
| ONT_UL | -cx ava-ont | **654522** | 682884 | **27:41:54** | 29:21:05 | 25.0 GB | **19.2 GB** |
| PacBio HiFi | -cx map-hifi | **274260** | 318231 | **9:53:33** | 11:08:45 | **12.9 GB** | 16.1 GB |
| ONT_RNA | -cx splice -uf -k14 | **121612** | 131063 | **4:23:17** | 4:37:06 | **21.9 GB** | 22.5 GB |
| BGI PCR-free | -ax sr | 432217 | **307897** | 15:16:59 | **12:33:02** | 14.9 GB | **12.6 GB** |
| Illumina PCR-free | -ax sr | 355364 | **306620** | 12:44:05 | **11:02:44** | 15.0 GB | **12.9 GB** |
| PacBio Onso | -ax sr | 161168 | **152442** | 5:44:25 | **5:20:48** | 14.9 GB | **12.5 GB** |
| AVITI 1000 bp insert | -ax sr | 620332 | **430574** | 21:45:35 | **17:12:46** | 14.9 GB | **12.6 GB** |
| AVITI 500–600 bp insert | -ax sr | 760693 | **608294** | 27:21:27 | **27:05:20** | 14.9 GB | **12.5 GB** |
| Illumina HiC | -ax sr --frag=no | 2410520 | **1727916** | 83:41:55 | **64:32:11** | 15.6 GB | **13.0 GB** |
| GRCh38 / GRCm38 | -x asm20 | **1522** | 2004 | **0:08:13** | 0:10:15 | 23.5 GB | **21.0 GB** |
| GRCh38 / T2T | -x asm5 | **6894** | 7084 | **0:54:06** | 1:00:54 | 30.3 GB | **28.2 GB** |

Different mapping workloads stress different stages of the seed-chain-extend pipeline, and the relative speedup of rammap over minimap2 in Table 2 reflects which stages rammap’s optimizations target most directly. Long-read overlap mapping (`-x ava-ont`) generates many seed hits per read pair and is dominated by chaining throughput; rammap’s vectorized chaining kernels (AVX2 on x86_64 and NEON on aarch64) produce a relative speedup on this workload. Reference mapping of error-prone long reads (`-cx map-ont`) similarly stresses chaining but additionally invokes full base-level DP to produce CIGAR strings, benefitting from both vectorized chaining and rammap’s wider DP SIMD targets. High-accuracy long-read mapping (`-cx map-hifi`) shifts the balance toward DP, since fewer, more confident chains are extended through longer base-level alignments; here the AVX512 DP path is the primary determinant of throughput on x86_64. By contrast, short-read (`-ax sr`) workloads spend a comparatively larger fraction of time in seed extraction, query batching, and output formatting— stages that are less amenable to SIMD acceleration in the current rammap implementation, and where minimap2 retains a performance advantage.

For short read presets, increased SIMD bandwidth is not beneficial and can cause overhead or throttling penalties on some architectures. For these large `sr` workloads, performance is limited by memory management and allocation overhead in chaining and DP, where Rust’s memory safety architecture is limiting compared to C. Empirically, alternative global memory allocators including `mimalloc` and `jemalloc` decrease runtime by a factor of 10-20% for accurate short reads, but increase maximum RSS by 10-20%. These may be selected at compile time (`--features [mi|je]malloc`), otherwise rammap uses Rust’s default system allocator.

## Conclusion

Rammap is a Rust reimplementation of minimap2 that produces identical output across all tested mapping presets, matches or exceeds wall-clock performance on long-read workloads, and confines memory-unsafe operations to its SIMD kernels with a fully scalar fallback for applications requiring provable safety end-to-end. Both the full mapping pipeline and the underlying base-level alignment routines are exposed as a library, and the modular pipeline admits substitution of seeding, chaining, and extension components—providing a practical foundation for embedding SIMD-accelerated mapping into domain-specific workflows within a memory-safe runtime.

## Code and Data Availability

Rammap is publicly available under an MIT license (https://github.com/jwanglab/rammap).

